# Genome-wide identification and analysis of A-to-I RNA editing events in the malignantly transformed cell lines from BEP2D induced by α-particles radiation

**DOI:** 10.1101/551499

**Authors:** Qiaowei Liu, Hao Li, Lukuan You, Tao Li, Lingling Li, Pingkun Zhou, Xiaochen Bo, Hebing Chen, Xiaohua Chen, Yi Hu

**Affiliations:** Medical School of Chinese PLA, Beijing 100853, P.R. China; Department of Medical Oncology, Chinese PLA General Hospital, Beijing 100853, P.R. China; Beijing Institute of Radiation Medicine, Beijing 100850, P.R. China

**Author notes:** Correspondence (Y.H.); (X.C.); (H.C.). These authors contributed equally to this work (Qiaowei Liu, Hao Li).

**Keywords:** RNA editing, radiation-induced cancer, BEP2D, oncogenes

## Abstract

Adenosine (A) to inosine (I) RNA editing is the most prevalent RNA editing mechanism in humans and play critical roles in tumorigenesis. However, the effects of radiation on RNA editing and the mechanisms of radiation-induced cancer were poorly understood. Here, we analyzed human bronchial epithelial BEP2D cells and radiation-induced malignantly transformed cells with next generation sequencing. By performing an integrated analysis of A-to-I RNA editing, we found that genome-encoded single-nucleotide polymorphisms (SNPs) might induce the downregulation of ADAR2 enzymes, and further caused the abnormal occurrence of RNA editing in malignantly transformed cells. These editing events were significantly enriched in differentially expressed genes between normal cells and cancer cells. In addition, oncogenes *CTNNB1* and *FN1* were highly edited and significantly overexpressed in cancer cells, thus may be responsible for the lung cancer progression. Our work provides a systematic analysis of RNA editing from lung tumor specimens with high-throughput RNA sequencing and DNA sequencing. Moreover, these results demonstrate further evidence for RNA editing as an important tumorigenesis mechanism.

## Introduction

Lung cancer remains the leading cause of cancer death in both men and women, and radon exposure is the second most common cause of lung cancer after smoking [1]. However, the molecular mechanisms of radon-induced lung cancer remain unclear.

RNA editing is a post-transcriptional modification process, the deamination of adenosines (A) to inosines (I) is the prominent RNA editing event in humans, where ADAR enzymes convert A to I without affecting the DNA sequence identity [2]. Intriguingly, RNA editing plays an important role in tumorigenesis, such as recoding RNA editing of AZIN1 predisposes to hepatocellular carcinoma[3], RNA editing in RHOQ promotes invasion potential in colorectal cancer [4], and *GABRA3* editing suppresses breast cancer metastasis [5]. Recent study has shown that, in non-small cell lung cancer samples, as ADAR gene amplification, the DNA repair enzyme NEIL1 (K242R) will increase recoding [6]. However, there were limited studies to date in further exploring the characteristics of RNA editing in lung cancer. What’s more, the effect of radiation on RNA editing is poorly understood.

Here, we investigate A-to-I RNA editing in human bronchial epithelial cells (BEP2D) and malignantly transformed cells (BERP35T1 and BERP35T4), which are important models to characterize the radiation-mediated carcinogenesis of lung [7,8]. By performing high-throughput RNA sequencing, we identified A-to-I editing sites with three robust bioinformatics methods. We then systemically compared editing events in normal cells and cancer cells. Further, by performing genome-wide DNA sequencing, we revealed that the genomic variants in *ADAR2* gene was responsible for the abnormal editing events in cancer cells. Finally, we reported two potential editing genes, *CTNNB1* and *FN1*, in lung cancer.

## Results

### Identification of A-to-I RNA editing

The prevalence and importance of A-to-I RNA editing have been illuminated in recent years largely owing to the rapid adoption of high-throughput sequencing technologies [9,10]. To analyze A-to-I RNA editing in BEP2D cells and malignantly transformed cell lines, we performed high-throughput RNA sequencing (RNA-Seq) on BEP2D cell and transformed BEP2D cells, which were irradiated with 1.5Gy dose of α-particles emitted by ^238^PuO_2_. Two transformed cell lines, BERP35T1 and BERP35T4, were investigated (Fig. 1A). Three biological replicates were sequenced and analyzed for each cell line. We then calculated gene expression level with Cufflinks program [11], for each cell line, biological replicates of RNA-seq showed highly reproducible results (Fig. 1B). Thus, our sequencing data were of high quality for downstream analysis.

**Figure. 1.**
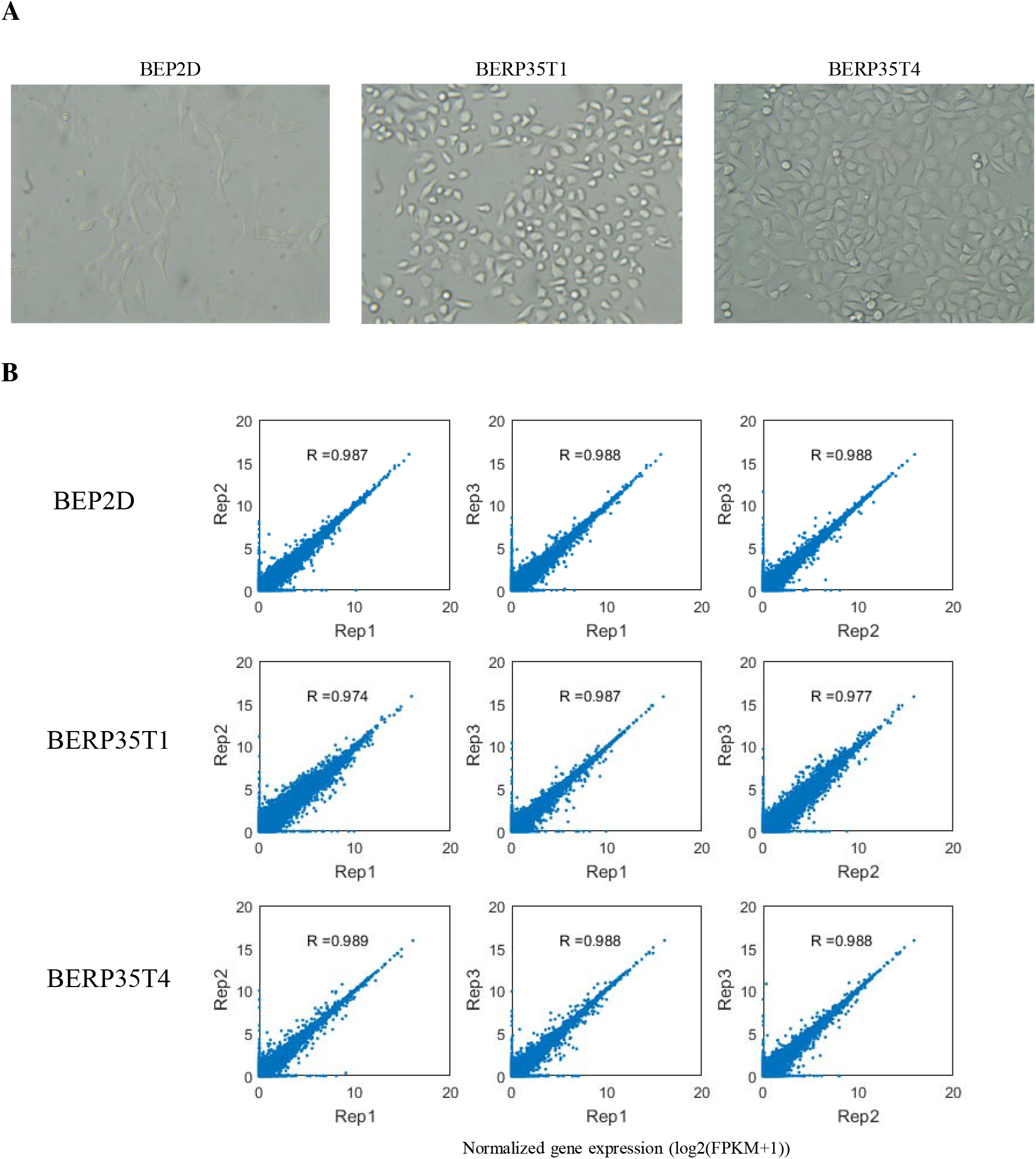
Cell culture and RNA sequencing. A. Photomicrographs of BEP2D (Left), BERP35T1 (Middle) and BERP35T4 (Right) showing the cellular atypia of the malignant transformed cells (100×). **B.** Scatter plots showing the consistency of normalized gene expression between biological replicates for each cell line.

Recent studies have reported that the most challenging part of identifying RNA editing is the discrimination of RNA editing sites from genome-encoded single-nucleotide polymorphisms (SNPs) and technical artifacts caused by sequencing or read-mapping errors [12-14]. To accurately identify RNA editing sites, we performed three famous methods including GIREMI [15], RNAEditor [16] and Separate method from Jin Billy Li [12] (See Methods). The GIREMI method combines statistical inference of mutual information (MI) between pairs of single-nucleotide variants (SNVs) in RNA-seq reads with machine learning to predict RNA editing sites. RNAEditor identify RNA editing by detecting ‘editing islands’. Separate method from Jin Billy Li identify RNA editing sites by strict filtering processes. For each sample, we only used RNA editing sites that can be detected in all three methods. For each cell line, we combined RNA editing sites from three biological replicates. As the most prevalent editing type in humans is adenosine-to-inosine (A-to-I) editing and most noncanonical editing are false positives [17], we only analyzed A-to-I RNA editing in this study. Final, 5659, 3820, 2446 A-to-I RNA editing sites were identified in BEP2D cell line and transformed cell lines BERP35T1 and BERP35T4, respectively (Table 1 and Supplemental Table 1).

**Table 1.**
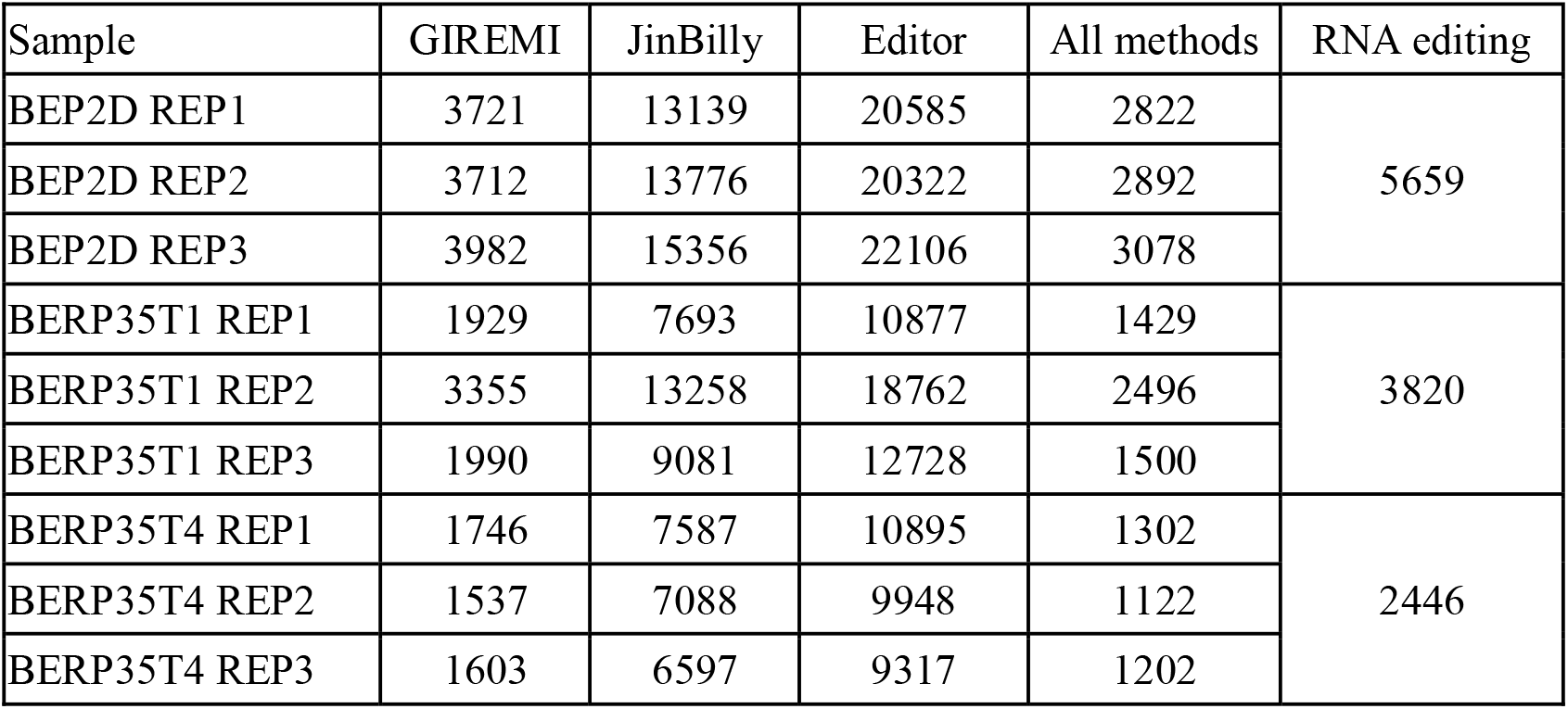
Summary of A-to-I RNA editing.

| Sample | GIREMI | JinBilly | Editor | All methods | RNA editing |
| --- | --- | --- | --- | --- | --- |
| BEP2D REP1 | 3721 | 13139 | 20585 | 2822 | 5659 |
| BEP2D REP2 | 3712 | 13776 | 20322 | 2892 |  |
| BEP2D REP3 | 3982 | 15356 | 22106 | 3078 |  |
| BERP35T1 REP1 | 1929 | 7693 | 10877 | 1429 | 3820 |
| BERP35T1 REP2 | 3355 | 13258 | 18762 | 2496 |  |
| BERP35T1 REP3 | 1990 | 9081 | 12728 | 1500 |  |
| BERP35T4 REP1 | 1746 | 7587 | 10895 | 1302 | 2446 |
| BERP35T4 REP2 | 1537 | 7088 | 9948 | 1122 |  |
| BERP35T4 REP3 | 1603 | 6597 | 9317 | 1202 |  |

### A-to-I RNA editing and associated genes in BEP2D and malignantly transformed cell

We next investigated the difference of A-to-I RNA editing between BEP2D cell line and malignantly transformed cell lines. First, 3,683∼4,217 editing events were disappeared and 1,004∼1,844 new editing events occurred from normal BEP2D cell to malignantly transformed cells, indicating a dramatic changes of RNA editing when BEP2D cell was irradiated (Fig. 2A). Generally, A-to-I editing is pervasive in Alu repeats because of the double-stranded RNA structures formed by inverted Alu repeats in many genes [18,19]. We found that although RNA editing is quite different in BEP2D and malignantly transformed cell lines, editing sites are still conserved in Alu repeats (Fig. 2B) and ∼60% of RNA editing events occurred in intergenic regions (Fig. 2C), thus, the basic distribution of A-to-I RNA editing did not change.

**Figure. 2.**
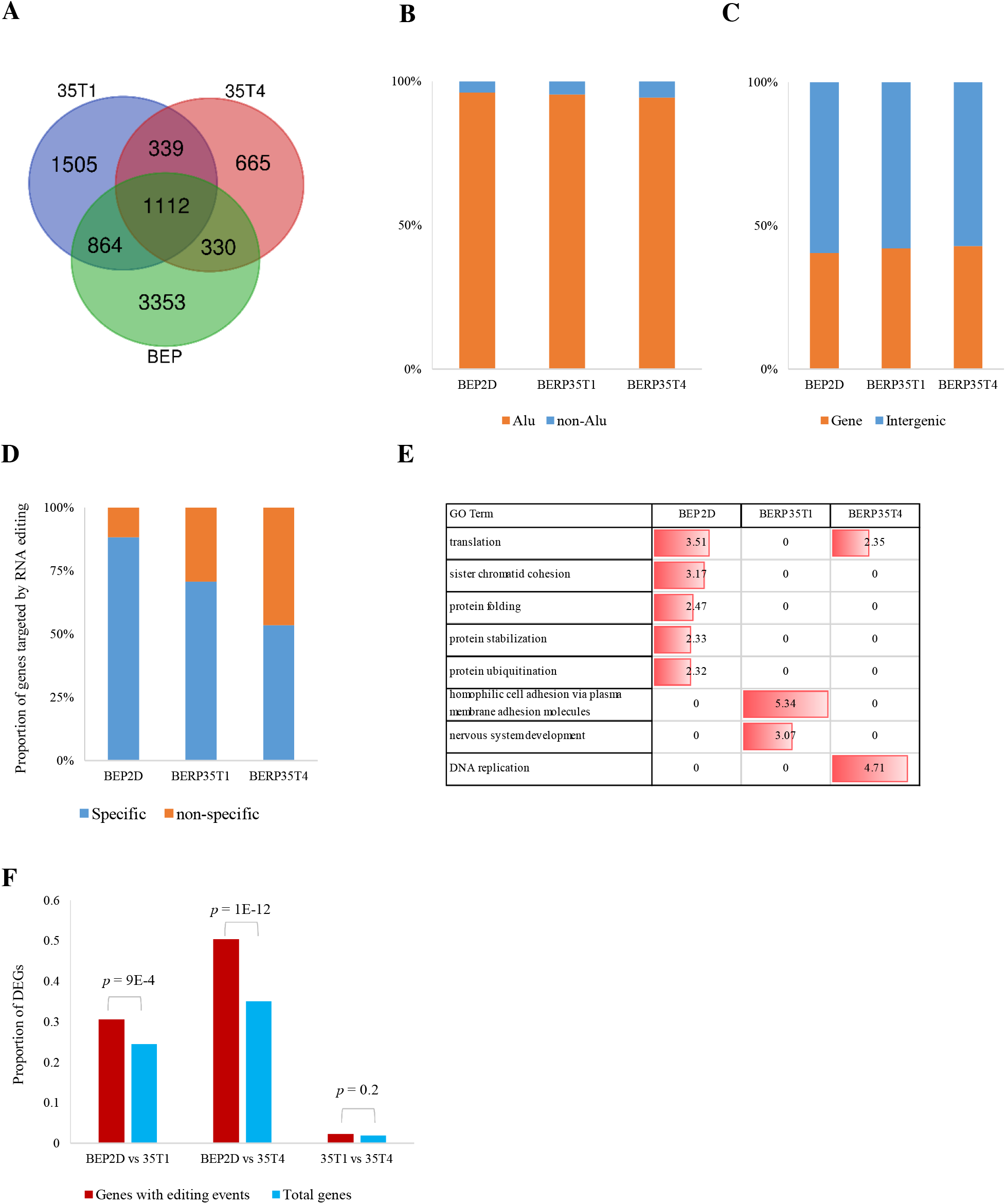
Identification and characterization of A-to-I RNA editing. **A.** Venn plot showing the overlaps of RNA editing sites in BEP2D, BERP35T1 and BERP35T4 cell lines. **B, C.** Bar plots showing the proportion of A-to-I RNA editing in Alu/non-Alu regions (B) and Genebody/Intergenic regions (C). **D.** Bar plot showing the proportion of tissue-specific editing genes in each cell line. **E.** GO enrichment for editing genes in each cell line, value was negative log10 of *p*-value. **F.** We compared the proportion of DEGs in total genes (blue bar) and DEGs in editing genes (red bar), *p*-value was calculated by hypergeometric test.

Next, we examined genes targeted by A-to-I RNA editing sites (editing genes for short). In general, 484, 426 and 305 genes were edited in BEP2D, BERP35T1 and BERP35T4 cell lines (Supplemental Table 2). We found that, in BEP2D cell line, 88% of editing genes were targeted by BEP2D-specific editing sites, but in BERP35T1 and BERP35T4, only 70% and 53% of editing genes were targeted by cell-specific editing sites (Fig. 2D). This result suggested that the editing rate of genes decreased when cell was irradiated and malignantly transformed. In addition, we performed gene ontology (GO) analysis to reveal the biological function of editing genes. We found that editing genes were enriched in different biological processes. For BEP2D, editing genes were enriched in protein processes and translation process, for BERP3T1, nervous system development and hemophilic cell adhesion process were highlighted and editing genes were enriched in DNA replication in BERP35T4 cell line (Fig. 2E).

To examine whether RNA editing affects transcription activity, we identified differentially expressed genes (DEGs) by performing Cuffdiff program [11] (Supplemental Table 3). We found that DEGs between normal BEP2D cell and malignantly transformed cells were significantly enriched in genes with RNA editing events (*p* value < 1E-3, hypergeometric test, Fig. 2F). This observation indicated that the dynamics of RNA editing sites was related to gene dysregulation and may further induce tumorigenesis.

### *ADAR2* down-regulation by genome SNPs

We next investigated the mechanism responsible for the differences observed in RNA editing between normal BEP2D cell and malignantly transformed cells. In human, A-to-I editing is performed by the ADAR family, which contains 3 genes: *ADAR1*, *ADAR2* and *ADAR3* [20-22]. We thus examined the transcript levels of *ADAR* genes. The expression level of *ADAR1* in BEP2D cell was comparable to that in BERP35T1 and BERP35T4 cell and *ADAR3* was silence in both BEP2D cell and malignantly transformed cells (Fig. 3A). However, *ADAR2* expression level significantly reduced in BERP35T1 and BERP35T4 (Fig. 3B). Previous studies have confirmed that *ADAR2* is lowly expressed in cancer e.g. glioblastoma [23], gastric cancer[24]. Thus the tumor progression of BEP2D seems to mainly be induced by *ADAR2* downregulation.

**Figure. 3.**
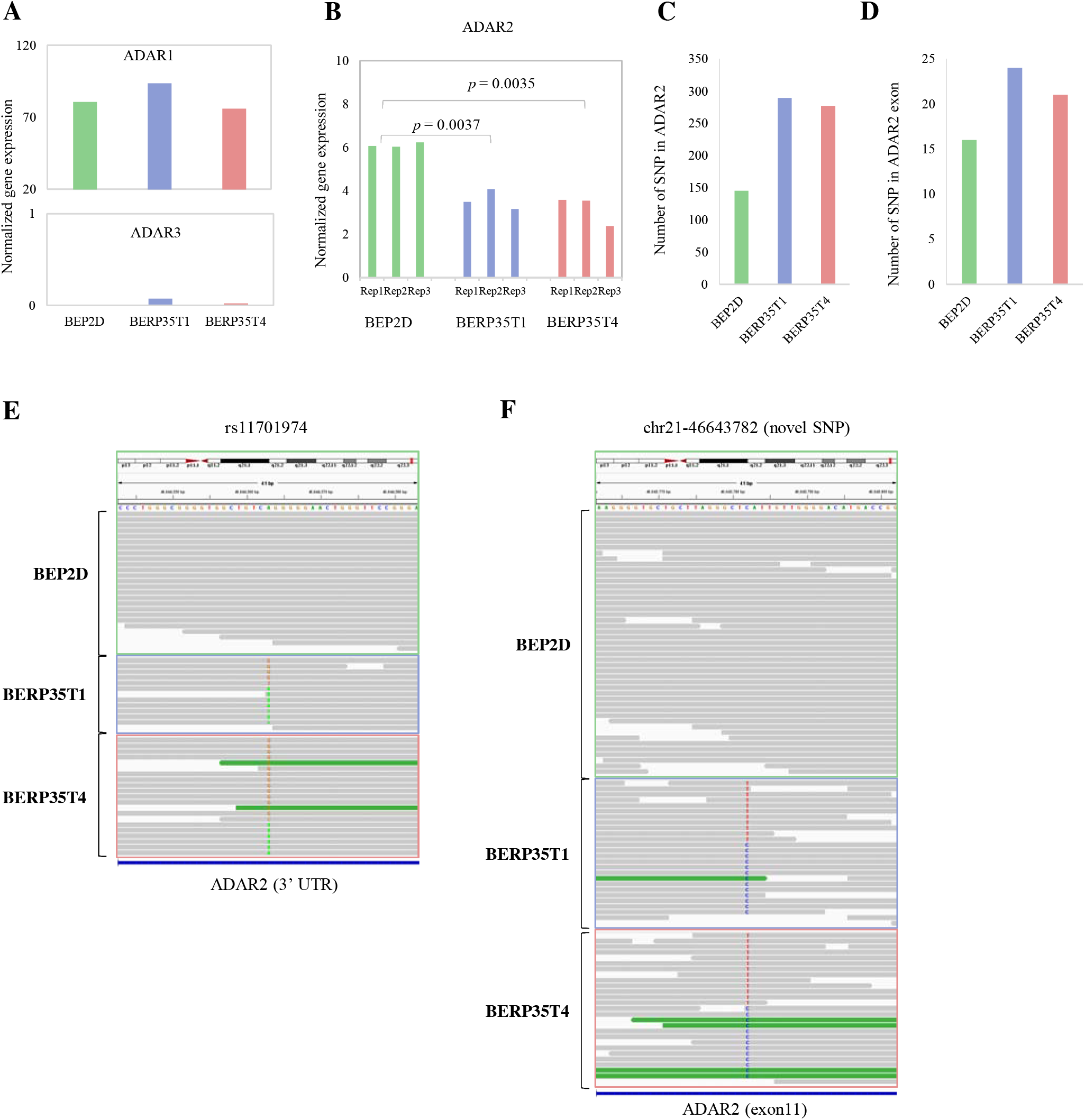
Down-regulation of ADAR2 induce RNA editing. **A, B.** Bar plots showing the normalized gene expression of *ADAR1, ADAR2* and *ADAR3* in each cell line. **C, D**. Bar plots showing the number of genomic variants in *ADAR2* (C) and exons of *ADAR2* (D). **E, F.** IGV plots showing the sequencing reads information of SNP rs11701974 and novel SNP chr21:46643782.

We further explored the possible mechanism of *ADAR2* downregulation in malignantly transformed cells. Radiation induced DNA alterations change gene expression and further increase cancer risk [25,26]. We next performed DNA sequencing (DNA-seq) on BEP2D cell line and malignantly transformed cells. We identified genome-encoded single-nucleotide polymorphisms (SNPs) using GATK pipeline [27] (See Methods). Surprisingly, nearly 2-fold SNPs in *ADAR2* gene were detected in malignantly transformed cells compared to normal BEP2D (Fig. 3C, Supplemental Table 4) and more SNPs in *ADAR2* exon were observed (Fig. 3D). For example, known SNP rs11701974, a genetic variant of HLA-DQB1 associated with human longevity [28], was detected in 3’ UTR of *ADAR2* and specific in BERP35T1 cell and BERP35T4 cell (Fig. 3E). Moreover, we identified 32 and 24 novel SNPs in BERP35T1 cell and BERP35T4 cell, respectively (Supplemental Table 4). For example, chr21:46643782 was altered in malignantly transformed cells (Fig. 3F). These results indicate that genomic variants induced by radiation may lead *ADAR2* downregulation and further decrease the editing rate of malignantly transformed cells.

### Oncogene *CTNNB1* and *FN1* are highly edited and significantly overexpressed in malignantly transformed cell

To gain insights into the biological relevance of RNA editing in malignantly transformed cell, we investigated 285 oncogenes from previous studies [29] (Supplemental Table 5). We found that oncogenes *CTNNB1, PABPC1* and *VHL* were edited in BERP35T1 cell. Notably, the expression level of *CTNNB1* in BERP35T1 cell was significantly higher than that in BEP2D cell (*p*-value = 0.00185, Fig. 4A). Previous studies reported that activating mutations in *CTNNB1* have oncogenic activity resulting in tumor development and somatic mutations are found in various tumor types [30-33]. We found two A-to-I editing events (chr3:41262966 and chr3:41262974) occurred in BERP35T1 cell and *CTNNB1* was overexpression in BERP35T1 cell (Fig. 4B). Similarly, we found three oncogenes *FN1*, *METTL14* and *VHL* were edited in BERP35T4 cell. Notably, the expression level of *FN1* in BERP35T4 cell was significantly higher than that in BEP2D cell (*p*-value = 5E-5, Fig. 4C). Previous studies have reported that transcriptional activation of *FN1* and gene fusions of *FN1* promote the malignant behavior of multiple cancers [34-36]. A strong A-to-I editing event (chr2:216236508) and a weak A-to-I editing event (chr2:216236482) were observed in BERP35T4 and *FN1* was overexpression in BERP35T4 cell (Fig. 4D). These results suggest that RNA editing are associated with oncogene overexpression and may further induce cancer progression.

**Figure. 4.**
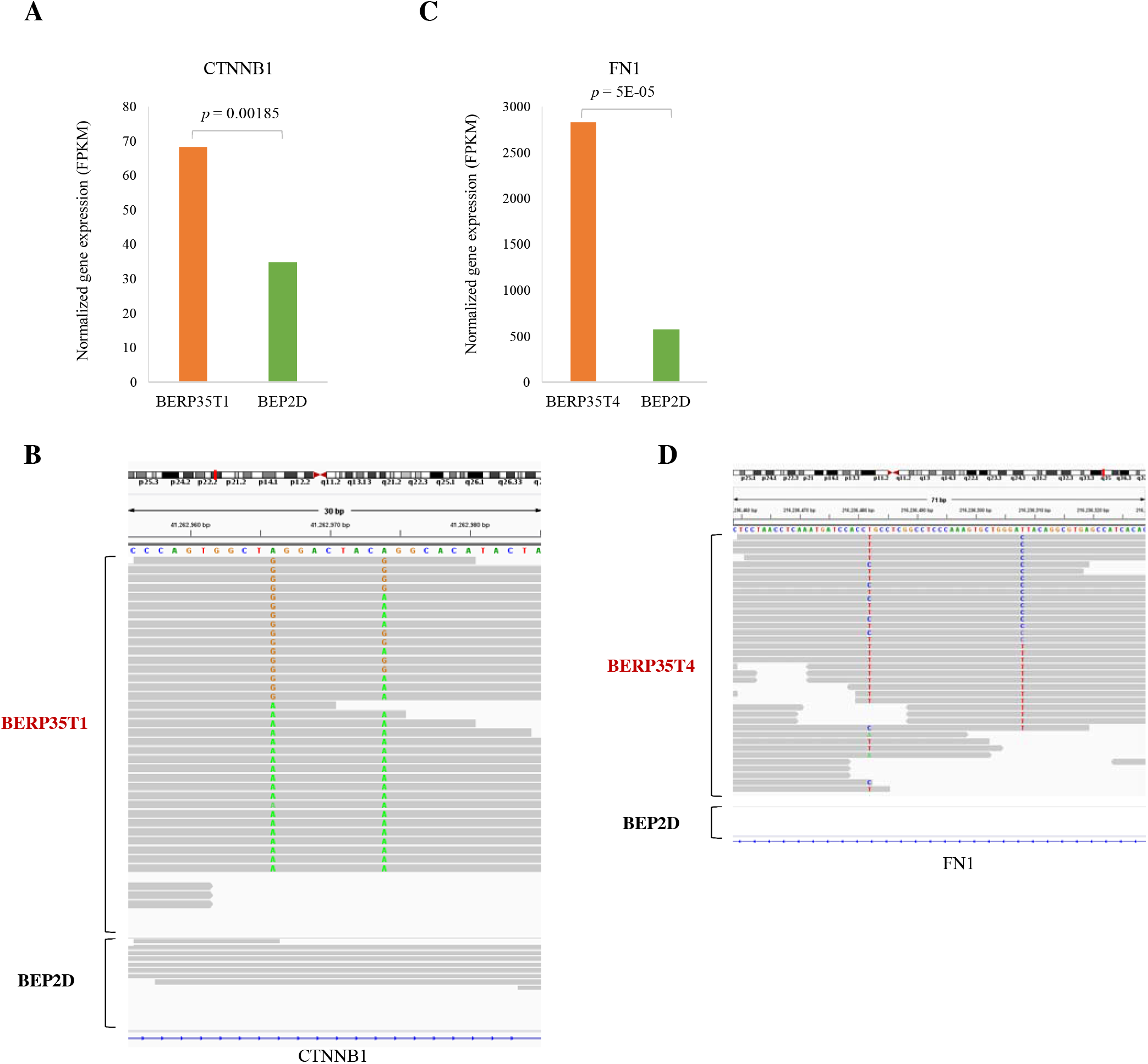
Oncogene CTNNB1 and FN1 are highly edited and significantly overexpressed in malignantly transformed cell. **A, C.** Bar plots showing the normalized gene expression of oncogenes *CTNNB1* and *FN1* in normal BEP2D cell and malignantly transformed BERP35T1 and BERP35T4 cells, respectively. **B, D.** IGV plots showing the editing events in oncogenes *CTNNB1* and *FN1*.

## Discussion

Radon is a recognized cause of lung cancer, however, the cellular and molecular mechanisms of radon-induced lung cancer remains unknown. To facilitate the study of this question, in our previous work, we established a model system of α-particle transformed human cells [37]. Then, we found a number of alterations in these cell models, including cytogenetics [38,39], gene expression [40], DNA repair [8], and genomic instability [41]. However, a genome-wide systematic analysis of this model based on next generation of sequencing is absent. In this work, we performed high-throughput RNA sequencing and genome-wide DNA sequencing to this model and discovered a new mechanism probably for tumorigenesis.

We all known the radon-oncogenic effect is DNA damage, but there were few articles about the effects of radiation on RNA editing. In this work, we provided genome-wide identification and analysis of A-to-I RNA editing events in the malignantly transformed cell line BEP2D induced by α-particles radiation, the results show that RNA editing sites changed greatly and the total amount decreased after radiation.

In cancer research, mutated proteins have been widely used as biomarkers and therapeutic targets, it is a fundamental issue to understanding the mechanisms contributing to protein diversity in cancer cells. RNA editing plays an important role in post-translational modification which can affect mRNA’s structure and stability [40,42], but little is known about how RNA editing operates in cancer [43]. Our work found that, in these cell models, genome-encoded single-nucleotide polymorphisms (SNPs) induced the downregulation of ADAR2 enzymes, and further caused the abnormal occurrence of RNA editing in malignantly transformed cells. Then, the abnormal occurrence of RNA editing led to abnormal expression of oncogenes, such as, *CTNNB1* and *FN1*, thus may be responsible for the lung cancer progression. These results demonstrate further evidence for RNA editing as an important tumorigenesis mechanism, and RNA editing sites might represent a new class of therapeutic targets.

## Materials and Methods

### Cell culture

The BEP2D cell line is a human papilloma-virus (HPV18)-immortalized human bronchial epithelial cell line and was established by Dr Curtis C. Harris (National Cancer Institute, MD, USA) [44]. These cells are anchorage dependent and non-tumorigenic in late passages. We got the authorization for research use only from Dr Curtis C. Harris and the Passage 20 of the BEP2D cell line was kindly provided by Tom K Hei (Center for Radiological Research, College of Physician and Surgeons, Columbia University, New York, USA) in the summer of 1993. The authentication of BEP2D cell line was tested by using short tandem repeats (STRs) analysis in June 2018. Although the information of BEP2D cells was not found in DSMZ and ATCC, our STRs results did match BBM cell line (ATCC Cell No. CRL-9482), BZR cell line (ATCC Cell No. CRL-9483) and BEAS-2B cell line (ATCC Cell No. CRL-9609), which are three tranformants derived from human bronchial epithelial cell (supplementary S1). The BERP35T1 and BERP35T4 malignant transformant cell lines were derived from BEP2D cells irradiated with 1.5 Gy of α-particle emitted from ^238^Pu source and were described in detail in a previous paper[8]. The cells were cultured in serum-free LHC-8 medium (Gibco, USA) at 37°C under a 95% air/5% CO2 atmosphere.

### RNA sequencing

Total RNAs were extracted from cells with RNAiso Reagent (TaKaRa, Dalian, China) following the manufacturer’s instruction. RNA degradation and contamination was monitored on 1% agarose gels. RNA purity was checked using the NanoPhotometer® spectrophotometer (IMPLEN, CA, USA). RNA concentration was measured using Qubit® RNA Assay Kit in Qubit® 2.0 Flurometer (Life Technologies, CA, USA). RNA integrity was assessed using the RNA Nano 6000 Assay Kit of the Bioanalyzer 2100 system (Agilent Technologies, CA, USA). A total amount of 1 µg RNA per sample was used as input material for the RNA sample preparations. Sequencing libraries were generated using NEBNext® Ultra™ RNA Library Prep Kit for Illumina® (NEB, USA) following manufacturer’s recommendations and index codes were added to attribute sequences to each sample. Briefly, mRNA was purified from total RNA using poly-T oligo-attached magnetic beads. Fragmentation was carried out using divalent cations under elevated temperature in NEBNext First Strand Synthesis Reaction Buffer (5X). First strand cDNA was synthesized using random hexamer primer and M-MuLV Reverse Transcriptase (RNase H-). Second strand cDNA synthesis was subsequently performed using DNA Polymerase I and RNase H. Remaining overhangs were converted into blunt ends via exonuclease/polymerase activities. After adenylation of 3’ ends of DNA fragments, NEBNext Adaptor with hairpin loop structure were ligated to prepare for hybridization. In order to select cDNA fragments of preferentially 150∼200 bp in length, the library fragments were purified with AMPure XP system (Beckman Coulter, Beverly, USA). Then 3 µl USER Enzyme (NEB, USA) was used with size-selected, adaptor-ligated cDNA at 37°C for 15 min followed by 5 min at 95 °C before PCR. Then PCR was performed with Phusion High-Fidelity DNA polymerase, Universal PCR primers and Index (X) Primer. At last, PCR products were purified (AMPure XP system) and library quality was assessed on the Agilent Bioanalyzer 2100 system.

The clustering of the index-coded samples was performed on a cBot Cluster Generation System using TruSeq PE Cluster Kit v3-cBot-HS (Illumia) according to the manufacturer’s instructions. After cluster generation, the library preparations were sequenced on an Illumina Hiseq platform and 150 bp paired-end reads were generated.

### DNA sequencing

Total DNAs were extracted from cells with DNAiso Reagent (TaKaRa, Dalian, China) following the manufacturer’s instruction. The quality of isolated genomic DNA was verified by using these two methods in combination: (1) DNA degradation and contamination were monitored on 1% agarose gels. (2) DNA concentration was measured by Qubit® DNA Assay Kit in Qubit® 2.0 Flurometer(Life Technologies, CA, USA). A total amount of 1µg DNA per sample was used as input material for the DNA library preparations. Sequencing library was generated using Truseq Nano DNA HT Sample Prep Kit (Illumina, USA) following manufacturer’s recommendations and index codes were added to each sample. Briefly, genomic DNA sample was fragmented by sonication to a size of 350 bp. Then DNA fragments were endpolished, A-tailed, and ligated with the full-length adapter for Illumina sequencing, followed by further PCR amplification. After PCR products were purified (AMPure XP system), libraries were analyzed for size distribution by Agilent 2100 Bioanalyzer and quantified by real-time PCR (3nM).

The clustering of the index-coded samples was performed on a cBot Cluster Generation System using Hiseq PE Cluster Kit (Illumina) according to the manufacturer’s instructions. After cluster generation, the DNA libraries were sequenced on Illumina Hiseq platform and 150 bp paired-end reads were generated.

### RNA editing identification

We adopted three previously published methods to accurately identify A-to-I RNA editing sites.

For Jinbilly’s method [12], we used the Burrows-Wheeler algorithm (BWA)[45] to align RNA-seq reads to a combination of the reference genome (hg19) and exonic sequences surrounding known splice junctions from available gene models. We chose the length of the splice junction regions to be slightly shorter than the RNA-seq reads to prevent redundant hits. Picard (http://picard.sourceforge.net/) was then used to remove identical reads (PCR duplicates) that mapped to the same location. GATK tools were used to perform local realignment around insertion and/or deletion polymorphisms and to recalibrate base quality scores. Variant calling was performed using GATK UnifiedGenotyper tool with options stand_call_conf of 0 and stand_emit_conf of 0. Further filtering were performed as described [46].

For GIREMI method [15], RNA-seq mapping and preprocessing was same as Jinbilly’s method. For each mismatch position, a total read coverage of ≧5 was required and the variant allele was required to be present in at least three reads. We then the following types of mismatches: those located in simple repeats regions or homopolymer runs of ≧ 5 nt, those associated with reads substantially biased toward one strand, those with extreme variant allele frequencies (>95% or <10%) and those located within 4 nt of a known spliced junction. Finally, we perform GIREMI tool to call RNA editing.

For RNAEditor [16], fastq format files from RNA-seq data were directly used as input for RNAEditor tools to call RNA editing sites.

### SNP identification

We used the Bowtie2 [47] to align DNA-seq reads to reference genome hg19, Picard (http://picard.sourceforge.net/) was then used to remove identical reads (PCR duplicates) that mapped to the same location. GATK tools were used to perform local realignment around insertion and/or deletion polymorphisms and to recalibrate base quality scores. Variant calling was performed using GATK UnifiedGenotyper tool.

### Statistical analysis

Gene expression was calculated using Cufflinks program default parameters. Differentially expressed genes (DEGs) were identified by Cuffdiff program, three biological replicates for each cell line were combined as the input of Cuffdiff and a *p*-value was reported to show the significance of DEGs. RNA editing targeted genes were assigned with ‘bedops’ program. GO analysis were performed by using DAVID [48].

### Accession numbers

The sequencing data have been deposited with the Gene Expression Omnibus under the accession ID GSE143212.

## Acknowledgements

We gratefully acknowledge funding from the Innovation Project of People’s Liberation Army (grant no.16CXZ042).

## Author Contributions

Conceived and designed the experiments: YH XC HC. Performed the experiments: QL HL. Analyzed the data: LY TL LL PZ XB. Contributed reagents/materials/analysis tools: QL HL HC. Wrote the paper: HL HC.

### Conflicts of Interest

The authors declare no conflict of interest.

## Supplementary Tables

Table S1. RNA editing sites in each cell line.

Table S2. Genes with RNA editing sites (sheet 1) and genes with tissue-specific RNA editing sites (sheet 2).

Table S3. Differentially expressed genes.

Table S4. SNPs in gene ADAR2 for each cell line.

Table S5. Oncogenes used in this study.

## Supplementary Materials

Report of Cell Line Authentication.

## References

1. Ferri G, Intranuovo G, Cavone D, Corrado V, Birtolo F, et al. (2018) Estimates of the Lung Cancer Cases Attributable to Radon in Municipalities of Two Apulia Provinces (Italy) and Assessment of Main Exposure Determinants. International Journal of Environmental Research & Public Health 15: 1294.

2. Bass BL (2002) RNA editing by adenosine deaminases that act on RNA. Annu Rev Biochem 71: 817–846.

3. Leilei C, Yan L, Chi Ho L, Chan THM, Chow RKK, et al. (2013) Recoding RNA editing of AZIN1 predisposes to hepatocellular carcinoma. Nature Medicine 19: 209–216.

4. Han SW, Kim HP, Shin JY, Jeong EG, Lee WC, et al. (2014) RNA editing in RHOQ promotes invasion potential in colorectal cancer. J Exp Med 211: 613–621.

5. Gumireddy K, Li A, Kossenkov AV, Sakurai M, Yan J, et al. (2016) The mRNA-edited form of GABRA3 suppresses GABRA3-mediated Akt activation and breast cancer metastasis. Nature Communications 7: 10715.

6. Anadón C, Guil S, Simóriudalbas L, Moutinho C, Setien F, et al. (2016) Gene amplification-associated overexpression of the RNA editing enzyme ADAR1 enhances human lung tumorigenesis. Oncogene 35: 4407–4413.

7. Liu C, Li P, Gao D, Zhou P, Shao Y, et al. (2014) Abnormal promoter methylation of multiple tumor suppressor genes in human bronchial epithelial malignant cells. Biomed Rep 2: 525–528.

8. Sun JF, Sui JL, Zhou PK, Geng Y, Hu YC, et al. (2002) Decreased efficiency of gamma-ray-induced DNA double-strand break rejoining in malignant transformants of human bronchial epithelial cells generated by alpha-particle exposure. Int J Radiat Biol 78: 773–780.

9. Gokul R, Billy LJ (2014) RADAR: a rigorously annotated database of A-to-I RNA editing. Nucleic Acids Research 42: D109.

10. Ramaswami G, Jin BL (2016) Identification of human RNA editing sites: A historical perspective. Methods 107: 42–47.

11. Trapnell C, Roberts A, Goff L, Pertea G, Kim D, et al. (2012) Differential gene and transcript expression analysis of RNA-seq experiments with TopHat and Cufflinks. Nat Protoc 7: 562–578.

12. Ramaswami G, Zhang R, Piskol R, Keegan LP, Deng P, et al. (2013) Identifying RNA editing sites using RNA sequencing data alone. Nat Methods 10: 128–132.

13. Kleinman CL, Adoue V, Majewski J (2012) RNA editing of protein sequences: a rare event in human transcriptomes. Rna 18: 1586–1596.

14. Kleinman CL, Majewski J (2012) Comment on “Widespread RNA and DNA sequence differences in the human transcriptome”. Science 335: 1302; author reply 1302.

15. Zhang Q, Xiao X (2015) Genome sequence–independent identification of RNA editing sites. Nature Methods 12: 347.

16. John D, Weirick T, Dimmeler S, Uchida S (2017) RNAEditor: easy detection of RNA editing events and the introduction of editing islands. Brief Bioinform 18: 993–1001.

17. Nishikura K (2010) Functions and regulation of RNA editing by ADAR deaminases. Annu Rev Biochem 79: 321–349.

18. Kim U, Wang Y, Sanford T, Zeng Y, Nishikura K (1994) Molecular cloning of cDNA for double-stranded RNA adenosine deaminase, a candidate enzyme for nuclear RNA editing. Proc Natl Acad Sci U S A 91: 11457–11461.

19. Levanon EY, Eisenberg E, Yelin R, Nemzer S, Hallegger M, et al. (2004) Systematic identification of abundant A-to-I editing sites in the human transcriptome. Nat Biotechnol 22: 1001–1005.

20. Higuchi M, Maas S, Single FN, Hartner J, Rozov A, et al. (2000) Point mutation in an AMPA receptor gene rescues lethality in mice deficient in the RNA-editing enzyme ADAR2. Nature 406: 78–81.

21. Lehmann KA, Bass BL (2000) Double-Stranded RNA Adenosine Deaminases ADAR1 and ADAR2 Have Overlapping Specificities. Biochemistry 39: 12875–12884.

22. Melcher T, Maas S, Herb A, Sprengel R, Seeburg PH, et al. (1996) A mammalian RNA editing enzyme. Nature 379: 460–464.

23. Maas S, Patt S, Schrey M, Rich A (2001) Underediting of glutamate receptor GluR-B mRNA in malignant gliomas. Proc Natl Acad Sci U S A 98: 14687–14692.

24. Chan THM, Qamra A, Tan KT, Jing G, Yang H, et al. (2016) ADAR-mediated RNA editing predicts progression and prognosis of Gastric Cancer. Gastroenterology 151: 637–650.e610.

25. Nath N, Esche J, Muller J, Jensen LR, Port M, et al. (2018) Exome Sequencing Discloses Ionizing-radiation-induced DNA Variants in the Genome of Human Gingiva Fibroblasts. Health Phys 115: 151–160.

26. Camero S, Ceccarelli S, De Felice F, Marampon F, Mannarino O, et al. (2018) PARP inhibitors affect growth, survival and radiation susceptibility of human alveolar and embryonal rhabdomyosarcoma cell lines. J Cancer Res Clin Oncol.

27. DePristo MA, Banks E, Poplin R, Garimella KV, Maguire JR, et al. (2011) A framework for variation discovery and genotyping using next-generation DNA sequencing data. Nat Genet 43: 491–498.

28. Yang F, Sun L, Zhu X, Han J, Zeng Y, et al. (2017) Correction: Identification of new genetic variants of HLA-DQB1 associated with human longevity and lipid homeostasis-a cross-sectional study in a Chinese population. Aging 9: 2316–2333.

29. Schroeder MP, Carlota RP, David T, Abel GP, Nuria LB (2014) OncodriveROLE classifies cancer driver genes in loss of function and activating mode of action. Bioinformatics 30: i549.

30. Hongbing L, Gibanananda R, Byong Hoon Y, Mete E, Rosen KV (2009) Down-regulation of death-associated protein kinase-2 is required for beta-catenin-induced anoikis resistance of malignant epithelial cells. Journal of Biological Chemistry 284: 2012.

31. J?Rg W, Frederik AK, Otmar H (2007) The tumor suppressor Fhit acts as a repressor of beta-catenin transcriptional activity. Proceedings of the National Academy of Sciences of the United States of America 104: 20344–20349.

32. Genovese G, Ghosh P, Li H, Rettino A, Sioletic S, et al. (2012) The tumor suppressor HINT1 regulates MITF and Î²-catenin transcriptional activity in melanoma cells. Cell Cycle 11: 2206–2215.

33. Reiko S, Miki S, Takafumi J, Kiyoko F, Kazufumi H, et al. (2012) β-catenin inhibits promyelocytic leukemia protein tumor suppressor function in colorectal cancer cells. Gastroenterology 142: 572–581

34. Wu J, Wang Y, Xu X, Cao H, Sahengbieke S, et al. (2016) Transcriptional activation of FN1 and IL11 by HMGA2 promotes the malignant behavior of colorectal cancer. Carcinogenesis 37: 511–521.

35. Lee JC, Su SY, Changou CA, Yang RS, Tsai KS, et al. (2016) Characterization of FN1-FGFR1 and novel FN1-FGF1 fusion genes in a large series of phosphaturic mesenchymal tumors. Mod Pathol 29: 1335–1346.

36. Zhan S, Li J, Wang T, Ge W (2018) Quantitative Proteomics Analysis of Sporadic Medullary Thyroid Cancer Reveals FN1 as a Potential Novel Candidate Prognostic Biomarker. Oncologist.

37. Sui JL, An J, Sun JF, Chen Y, Wu DC, et al. (2004) Spindle checkpoint and apoptotic response in α-particle transformed human bronchial epithelial cells. Radiation & Environmental Biophysics 43: 257–263.

38. Weaver DA, Hei TK, Hukku B, McRaven JA, Willey JC (1997) Cytogenetic and molecular genetic analysis of tumorigenic human bronchial epithelial cells induced by radon alpha particles. Carcinogenesis 18: 1251–1257.

39. Sui JL, Liu XL, Lou TZ, Shi_Li GE, Geng Y, et al. (2002) Cytogenetic analysis of malignant transformants of human bronchial epithelial cells BEP2D generated by α-particles exposure. Bulletin of the Academy of Military Medical Sciences 2: 86–89.

40. Zhao YL, Piao CQ, Hall EJ, Hei TK (2001) Mechanisms of radiation-induced neoplastic transformation of human bronchial epithelial cells. Radiation Research 155: 230–234.

41. Piao CQ, Hei TK (2001) Gene amplification and microsatellite instability induced in tumorigenic human bronchial epithelial cells by alpha particles and heavy ions. Radiation Research 155: 263.

42. Nishikura K (2010) Functions and regulation of RNA editing by ADAR deaminases. [Review] [155 refs]. Annual Review of Biochemistry 79: 321.

43. Fumagalli D, Gacquer D, Rothé F, Lefort A, Libert F, et al. (2015) Principles Governing A-to-I RNA Editing in the Breast Cancer Transcriptome. Cell Reports 13: 277–289.

44. Willey JC, Broussoud A, Sleemi A, Bennett WP, Cerutti P, et al. (1991) Immortalization of normal human bronchial epithelial cells by human papillomaviruses 16 or 18. Cancer Res 51: 5370–5377.

45. Li H, Durbin R (2010) Fast and accurate long-read alignment with Burrows-Wheeler transform. Bioinformatics 26: 589–595.

46. Ramaswami G, Lin W, Piskol R, Tan MH, Davis C, et al. (2012) Accurate identification of human Alu and non-Alu RNA editing sites. Nat Methods 9: 579–581.

47. Langmead B, Salzberg SL (2012) Fast gapped-read alignment with Bowtie 2. Nat Methods 9: 357–359.

48. Huang da W, Sherman BT, Lempicki RA (2009) Systematic and integrative analysis of large gene lists using DAVID bioinformatics resources. Nat Protoc 4: 44–57.

