## Supplemental Tables for "Genome-wide identification and analysis of A-to-I RNA editing events in the malignantly transformed cell lines from BEP2D induced by α-particles radiation": Supplementary Materials S1.pdf

### Report of Cell Line Authentication

**Service Code :** LWXB18023

**Report Date :** 2018.6.7

**BGI**

| Project Information | | | |
|---|---|---|---|
| Project No. | LWXB18023 | Project Name | BEP2D cell line authentication |
| Customer Name | Qiaowei Liu | Customer Unit | Beijing Institute of Radiation Medicine |
| Customer Contact | | PM | ShuangyuWang |
| Sale name | YamingChen | | |
| **Overview of Services** | | | |
| Testing Date | 2018.6.4 | Report Date | 2018.6.7 |
| Sample Number | 1 | | |
| Actual receivable sum | RMB800 | | |
| **Note** | | | |

### Catalog

### 1. Experiment Objectives

Cell information identification

### 2. Experimental procedure and method

#### 2.1 Experiment Reagents

HUMDNA TYPING(Yanhuang)

#### 2.2 Experiment Apparatus

GeneAmp® PCR system9700， ABI3730XL

#### 2.3 Experimental Procedure

PCR amplification system:

| | | |
|---|---|---|
| ddH2O | 2 | µl |
| STR21 2X Master Mix | 5 | µl |
| STR21 4X Primer Pair Mix | 2 | µl |
| DNA | 1 | µl |
| Total | 10 | µl |

PCR amplification reaction procedures: Applied Biosystems 9700 PCR System

| | | |
|---|---|---|
| 91℃ | 1 min | |
| 95℃ | 10 sec | ×28Cycle |
| 58℃ | 1min | ×28Cycle |
| 70℃ | 20 sec | ×28Cycle |
| 60℃ | 30 min | |
| 4℃ | ∞ | |

#### 2.4 Detection

Amplified products were separated using an Applied Biosystems® 3730XL Genetic Analyzer. Experimental Process ： Amplified products— Add Internal Lane Standard—template degeneration—Detection(Pre-electrophoresis: 1.2kV 5min, Electrophoresis :7.5kV 2h)—Analysis.

| | |
|---|---|
| STR-500 | 0.5µl |
| PCR products | 1µl |
| HIDI | 8.5µL |
| Total | 10µL |

### 3. Results and Data

#### 3.1 BEP2D-Lqw STR Data

| STR Loci | Results | |
|---|---|---|
| Yindel | 2 | |
| AMEL | X | Y |
| D3S1358 | 15 | 17 |
| D13S317 | 13 | |
| D7S820 | 10 | 13 |
| D16S539 | 12 | |
| SE33 | 20.2 | 28.2 |
| D10S1248 | 14 | 15 |
| D5S818 | 12 | 13 |
| D21S11 | 28 | 30 |
| TPOX | 6 | 11 |
| D1S1656 | 14 | |
| D6S1043 | 12 | 18 |
| DXS6795 | 11 | |
| D19S433 | 13.2 | 15.2 |
| D22S1045 | 14 | 16 |
| D8S1179 | 13 | 15 |
| Penta E | 5 | 8 |
| DYS391 | 10 | |
| D2S441 | 11 | 11.3 |
| D12S391 | 17 | 18 |
| D2S1338 | 22 | 23 |
| vWA | 17 | 18 |
| Penta D | 2.2 | 13 |
| TH01 | 7 | 9.3 |
| D18S51 | 18 | 19 |
| CSF1PO | 9 | 12 |
| FGA | 20 | 24 |

#### 3.2 BEP2D-Lqw STR Loci Data Comparison

Search result in DSMZ database:

| EV | Cell No. | Cell name | Locus names | | | | | | | | |
|---|---|---|---|---|---|---|---|---|---|---|---|
| | | | D5S818 | D13S317 | D7S820 | D16S539 | VWA | TH01 | AM | TPOX | CSF1PO |
| | Query (Your Cell) | | 12,13 | 13,13 | 10,13 | 12,12 | 17,18 | 7,9.3 | X,Y | 6,11 | 9,12 |
| 1.00(36/36) | CRL-9482 | BBM | 12,13 | 13,13 | 10,13 | 12,12 | 17,18 | 7,9.3 | X,Y | 6,11 | 9,12 |
| 1.00(36/36) | CRL-9483 | BZR | 12,13 | 13,13 | 10,13 | 12,12 | 17,18 | 7,9.3 | X,Y | 6,11 | 9,12 |
| 1.00(36/36) | CRL-9609 | BEAS-2B | 12,13 | 13,13 | 10,13 | 12,12 | 17,18 | 7,9.3 | X,Y | 6,11 | 9,12 |
| 0.67(24/36) | CRL-7065 | Hs 97.Fs | 11,12 | 12,13 | 10,13 | 12,12 | 16,16 | 7,9.3 | X,Y | 8,11 | 10,12 |
| 0.61(22/36) | CRL-11233 | THLE-3 | 13,13 | 13,13 | 8,10 | 11,12 | 17,18 | 8,9.3 | X,X | 6,9 | 11,12 |

BEP2D cells information was not found in DSMZ and ATCC.

Note: Reference standards: ANSI/ATCC, Authentication of Human Cell Line Standardization of STR Profiling. 2011, ASN - 0002-0002. STR results matches more than 80% can be considered the same source of cells.

#### 3.3 BEP2D-Lqw STR Profiles

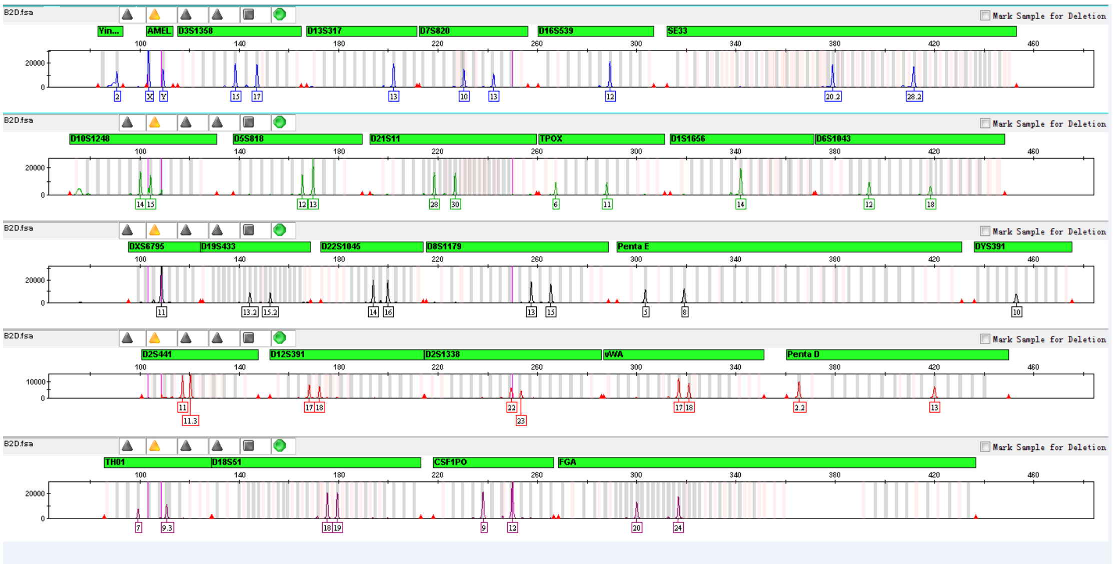

### 4. Reference

[1] Zhao, M., et al., Assembly and initial characterization of a panel of 85 genomically validated cell lines from diverse head and neck tumor sites. Clin Cancer Res, 2011. 17(23): p. 7248-64.

[2] Masters, J.R., Cell-line authentication: End the scandal of false cell lines. Nature, 2012. 492(7428): p. 186.

[3]American Type Culture Collection Standards Development Organization Workgroup, A.S.N., Authentication of Human Cell Lines: Standardization of STR Profiling. 2011, ANSI/ATCC ASN-0002-2011.

[4] Reid, Y.A., Characterization and authentication of cancer cell lines: an overview. Methods Mol

Biol, 2011. 731: p. 35-43.
[5] Lorsch, J.R., F.S. Collins, and J. Lippincott-Schwartz, Cell Biology. Fixing problems with cell lines. Science, 2014. 346(6216): p. 1452-3.
[6] Chatterjee, R., Cell biology. Cases of mistaken identity. Science, 2007. 315(5814): p. 928-31.
